# Network topology determines neuronal phase-of-firing codes

**DOI:** 10.64898/2026.07.23.740383

**Authors:** M. R. Pratyush, Collins Assisi

**Affiliations:** Indian Institute of Science Education and Research, Pune - 411008, India; Carnegie Mellon University, Pittsburgh, PA 15213, USA

## Abstract

Brain rhythms organize neural activity in time. When a shared oscillation is broadcast across a population of neurons, the phase at which the neurons fire with respect to the oscillation can itself carry information, a strategy the brain appears to use for navigation, memory, and sensory perception. In such a phase-of-firing code, information is conveyed not by how rapidly a neuron fires but by when it fires relative to the phase of the ongoing rhythm. Models that implement such phase-of-firing codes consider the neurons as independent. How such codes arise in cortical and hippocampal circuits, where neurons are densely interconnected rather than independent, remains unclear. Here we show that recurrent connectivity does not merely permit phase-of-firing codes but actively determines which patterns the network can hold onto. Using networks of model neurons wired in a manner determined by the constraints of a Sudoku puzzle, we find that the network supports an enormous repertoire of distinct rhythmic firing patterns, each corresponding to a valid Sudoku solution and selectable by brief targeted inputs against a common background oscillation. Some patterns persist after the input is removed while others dissolve. We show that a factor deciding this selectivity is network symmetry. Patterns that respect the symmetries of the wiring are stable, while those that break them are not, and small perturbations can switch the network between stable patterns. Adjusting synaptic weights to remove asymmetry stabilizes patterns that were previously unstable, giving the network a tunable memory. More broadly, our results suggest that a common oscillatory drive, targeted biases to the neurons, and recurrent network structure, act together as complementary control variables. The oscillation sets up a temporal scaffold, the biases select spatiotemporal patterns, and the connectivity determines the stability of the chosen patterns. Tuning these complementary variables can predictably and flexibly reshape the repertoire of activity a circuit expresses.

**Significance Statement:** Many brain regions generate rhythmic activity, and the precise moment at which individual neurons fire within each rhythm carries information that the brain can harness to encode memory, perception and location as an animal navigates its environment. In such a phase-of-firing code, information is conveyed not by how rapidly a neuron fires but by when it fires relative to the phase of the ongoing network rhythm. Existing theories of how phase-of-firing codes arise have modeled neurons as independent, leaving open how such codes emerge in the densely interconnected circuits of cortex and hippocampus. Using model networks wired to reflect the constraints of the Sudoku puzzle, we show that recurrent connectivity supports a vast repertoire of distinct rhythmic firing patterns, and that the symmetries of the wiring determines which of those patterns the network can stably hold. This identifies network symmetry as a general principle relating circuit architecture to temporal coding.

## Introduction

In neuronal networks, information can be encoded by which neurons fire and by their time-varying firing rates. When a common oscillation is broadcast across a network, an additional coding dimension becomes available, namely, the phase of the oscillation at which individual neurons spike. Because oscillatory activity in the brain spans a wide range of frequencies, from about 0.5 Hz to 500 Hz^1^, many brain networks are likely to exploit this **phase-of-firing code**^2^. For example, place cells in the rat hippocampus and grid cells in the entorhinal cortex fire at progressively earlier phases of a 4-10 Hz theta oscillation as the animal moves through a cell’s place field, so that the phase of firing encodes the animal’s precise position within that field^3–5^. This phase-of-firing code has additionally been implicated in higher sensory areas, including visual and auditory cortex (see, for instance, the examples summarized in Masquelier *et al*., 2009^6^).

How do phase-of-firing codes emerge in neuronal networks? Hopfield proposed that an analog variable, such as odor concentration or sound intensity, could be encoded in the timing of action potentials relative to a common oscillation^7^. In this scheme, the neuron receives both an oscillatory input and a constant depolarizing drive (**Fig. 1a**). As the strength of the depolarizing drive increases, the neuron fires its spike earlier within each cycle of the background oscillation (**Figs. 1a, b and S1**). The analog intensity is thus encoded in the phase of firing^7,8^. Extending this idea to a population of neurons yields a high-dimensional phase-of-firing code where different neurons receive different depolarizing inputs and spike at different phases, generating complex spatiotemporal activity patterns. However, existing models construct such codes using uncoupled neurons^7,8^, and thus largely ignore how network architecture shapes these phase relationships. To illustrate the problem posed by recurrent networks, consider the simple bi-partite network shown in **Fig. 1c**. The network consists of two groups of neurons with no within-group connectivity, but all-to-all inhibitory connections across groups. Both groups of neurons are driven by a common oscillatory drive and a uniform depolarizing input.

**Figure 1:**
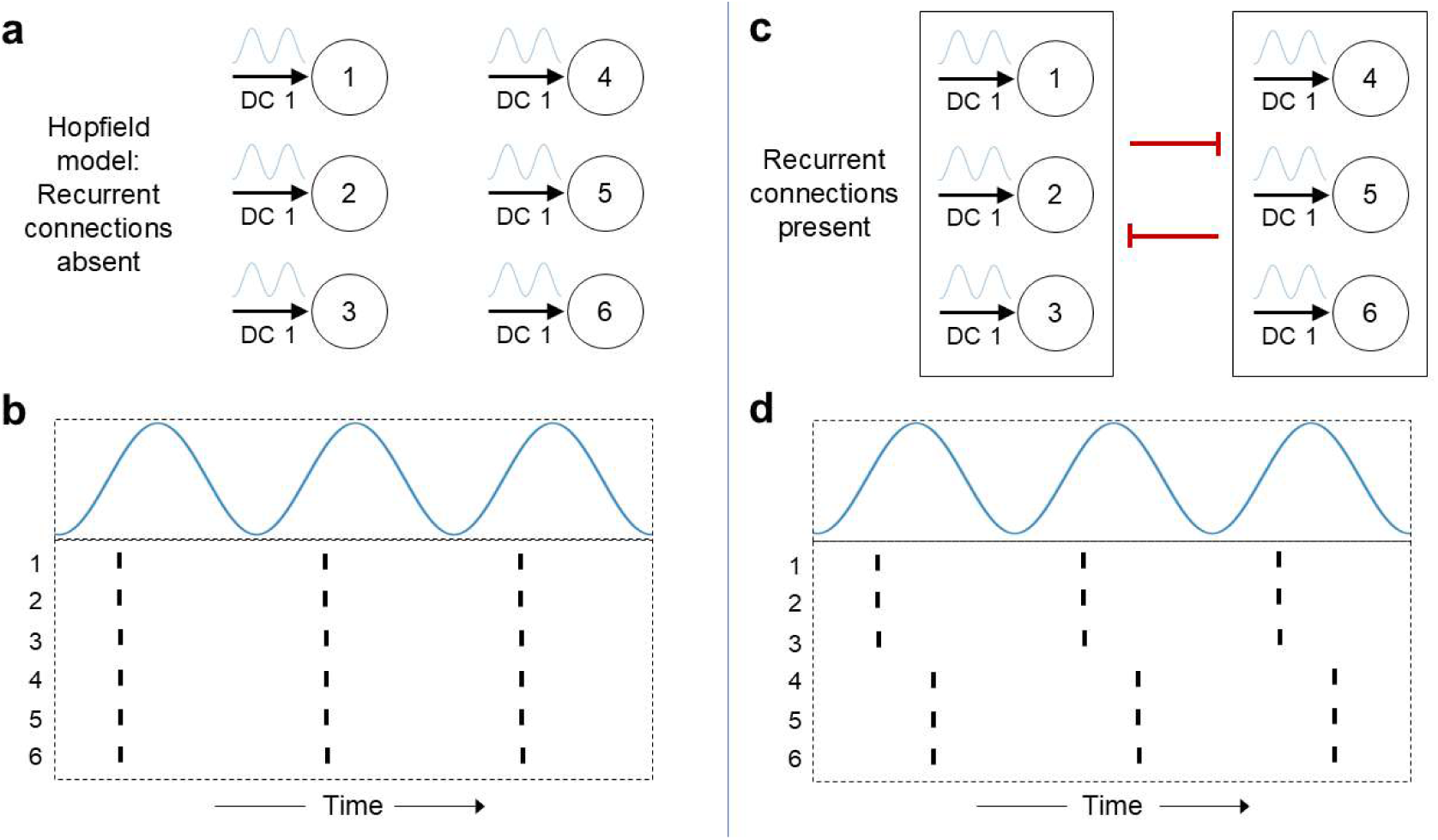
Recurrent inhibition disrupts synchronous spiking. (a–b) Uncoupled network neurons driven by a common subthreshold oscillation and common input bias (DC1) spike synchronously. (c–d) Adding recurrent inhibitory connections results in two groups of synchronously spiking neurons.

Unlike the uncoupled network (**Fig. 1a**) where all neurons spike synchronously at a phase determined by the input (**Fig. 1b**), neurons in this reciprocal inhibitory network fall into two synchronous groups (**Fig. 1d**) that spike at different phases of the oscillation despite receiving the same oscillatory drive and depolarization.

Here, we extend Hopfield’s model to implement a phase-of-firing code in recurrent excitatory-inhibitory networks. In these networks, we can embed multiple steady states and selectively access them with targeted depolarizing inputs. In many cases, depending on the symmetries of the network, the resulting states persist even after these inputs are removed. By contrast, when recurrent connections are eliminated, all neurons synchronize once the biasing input is turned off. Transient perturbations can also drive switches between distinct network states, whose activity patterns continue to persist after the perturbation decays. In sum, our results show that network topology determines which spatiotemporal patterns a circuit can stably express.

## Model

In our model, the key structural ingredient is competition in the inhibitory subnetwork. Neurons that reciprocally inhibit each other tend to spike at different times. The dynamics of such networks can be related to a graph-theoretic notion, coloring, in which neurons are nodes and inhibitory synapses are edges^9,10^. A coloring assigns different colors to any two neurons connected by an inhibitory edge. For example, the tripartite network in **Fig. 2a, b** can be minimally colored with three colors, and in this particular case only one minimal coloring exists. In this simple network, neurons assigned different colors spike at different times. In general, a network can admit many distinct colorings with each coloring corresponding to a different dynamical state. To obtain a rich set of such colorings, we construct a network based on the Sudoku puzzle^11,12^. Each solution of a Sudoku puzzle can be mapped to a coloring of this network^11^. As a concrete example, consider a 4×4 Sudoku consisting of a 4×4 grid in which the digits 1-4 must each appear exactly once in every row, column, and 2×2 sub-grid (**Fig. 2c, d**). To build the 4×4 Sudoku network, we place a neuron in each cell of an empty 4×4 grid and connect neurons within the same row, column, or 2×2 sub-grid by fast reciprocal inhibition (**Fig. 2c**). Each valid 4-coloring of this network then corresponds to a 4×4 Sudoku solution satisfying all constraints. In an earlier study, we showed that the dynamics of this network evolve so that neurons assigned the same color spike synchronously, while neurons with different colors spike at distinct times^12^. Thus, for a given set of initial conditions, the network activity converges to a specific Sudoku solution (i.e., one particular coloring). In a purely inhibitory network, neurons in different color classes tend to spike at distinct times, while neurons in the same class may spike together if they receive similar depolarizing input; however, within-group synchrony is not guaranteed because group members do not directly interact and are subject to noise and differing initial states. Excitatory synapses, in contrast, promote synchrony, so we add excitatory connections complementary to inhibition: any pair of neurons that is not reciprocally inhibitory is coupled by reciprocal excitation, with identical weights for all inhibitory synapses, and identical weights for all excitatory synapses (in general different from the inhibitory weights). By tuning the excitation-inhibition ratio, we identify a regime in which neurons within each color group spike synchronously and the temporal spiking pattern of the network can be directly mapped onto a coloring of the underlying graph. Under different initial conditions, the dynamics converge to different periodic states, each of which can be mapped to a distinct Sudoku solution (see Chowdhary and Assisi, 2019^12^ for details).

**Figure 2:**
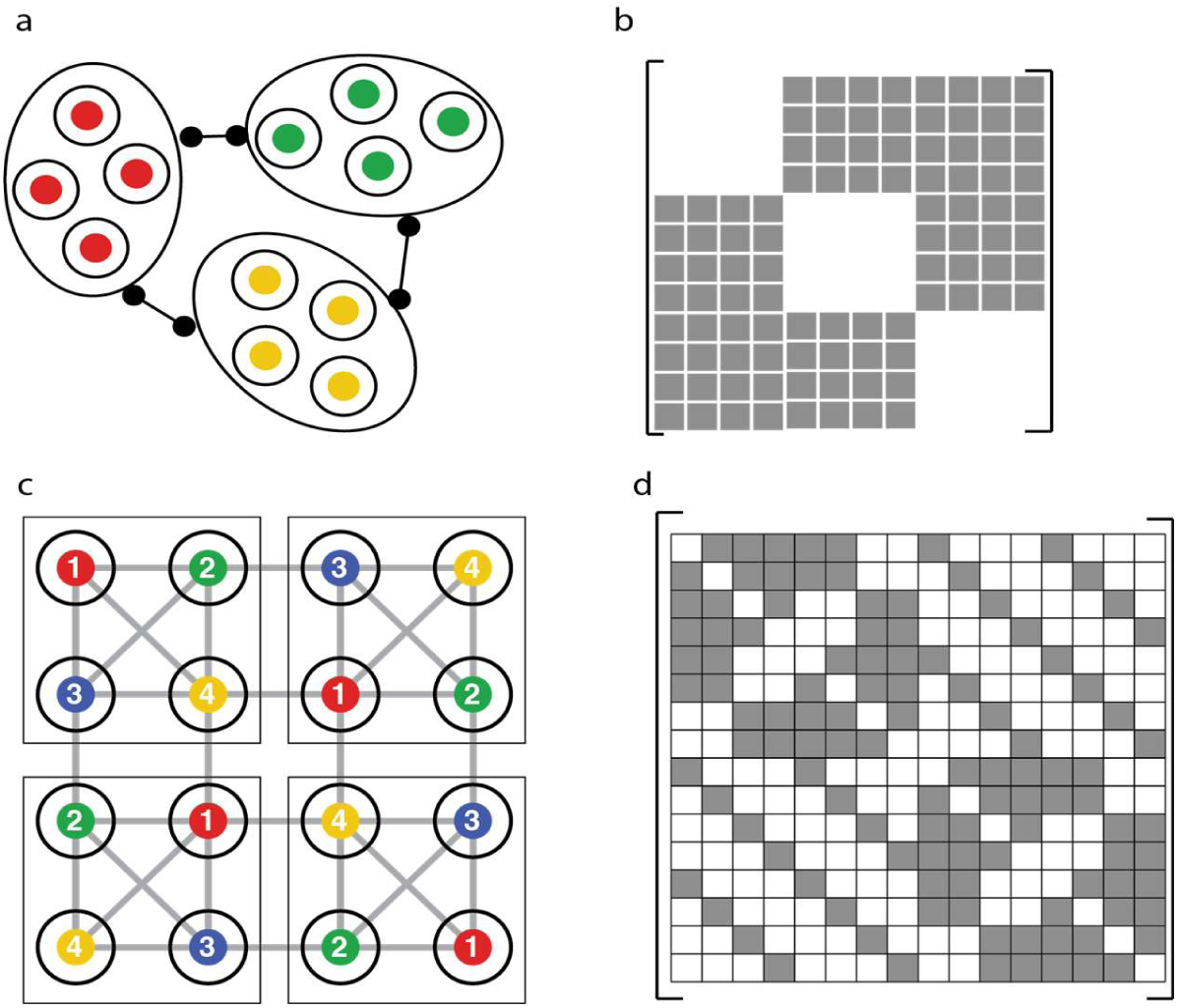
Network architectures and colorings. (a) Tripartite network with mutual inhibition, minimally colored with 3 colors. (b) Adjacency matrix of the network in (a). Shaded entries in the matrix are assigned 1. Other entries are set to 0. (c) 4×4 Sudoku network with all-to-all inhibition within rows, columns, and 2×2 subgrids. (d) Adjacency matrix of the network in (c).

## Results

### Using oscillations to access specific spatiotemporal patterns

In general, a network may admit several colorings, and thus multiple periodic attractor states. The spiking activity of a periodic state can be visualized as a raster plot (**Fig. 3a, left**) or a circular “phase” plot representing a single cycle (**Fig. 3b**). Starting with arbitrary initial conditions, the network typically displays transient aperiodic spiking activity before approaching an asymptotic state characterized by a periodic sequence of spikes (**Fig. 3b**). For a Sudoku network, the asymptotic periodic state corresponds to a coloring of the network. Neurons (nodes) that spike synchronously can be assigned the same color or digit, which can then be mapped back onto the Sudoku grid to yield a valid solution (**Fig. 3a, right**). Our simulations show that the asymptotic state depends on the initial condition (**Fig. 3c**). Because the number of attractors is large and the phase space is high-dimensional, mapping each basin of attraction can be intractable. We therefore asked whether we can reliably drive the network from an arbitrary initial state to a chosen attractor (i.e., a particular coloring or Sudoku solution). In the absence of inhibitory connections, neurons driven by a common oscillatory input spike synchronously at a fixed phase of the oscillation^8^ (**Fig. 1a, b**). A depolarizing bias to individual neurons can advance or retard the phase at which that neuron spikes. Over a range of bias currents, each neuron spikes exactly once per cycle, and its spiking phase varies monotonically with the bias (**Fig. S1**; Fig. 1 of Brody and Hopfield, 2003^8^). We restricted our simulations to this range of bias currents (see Methods).

**Figure 3:**
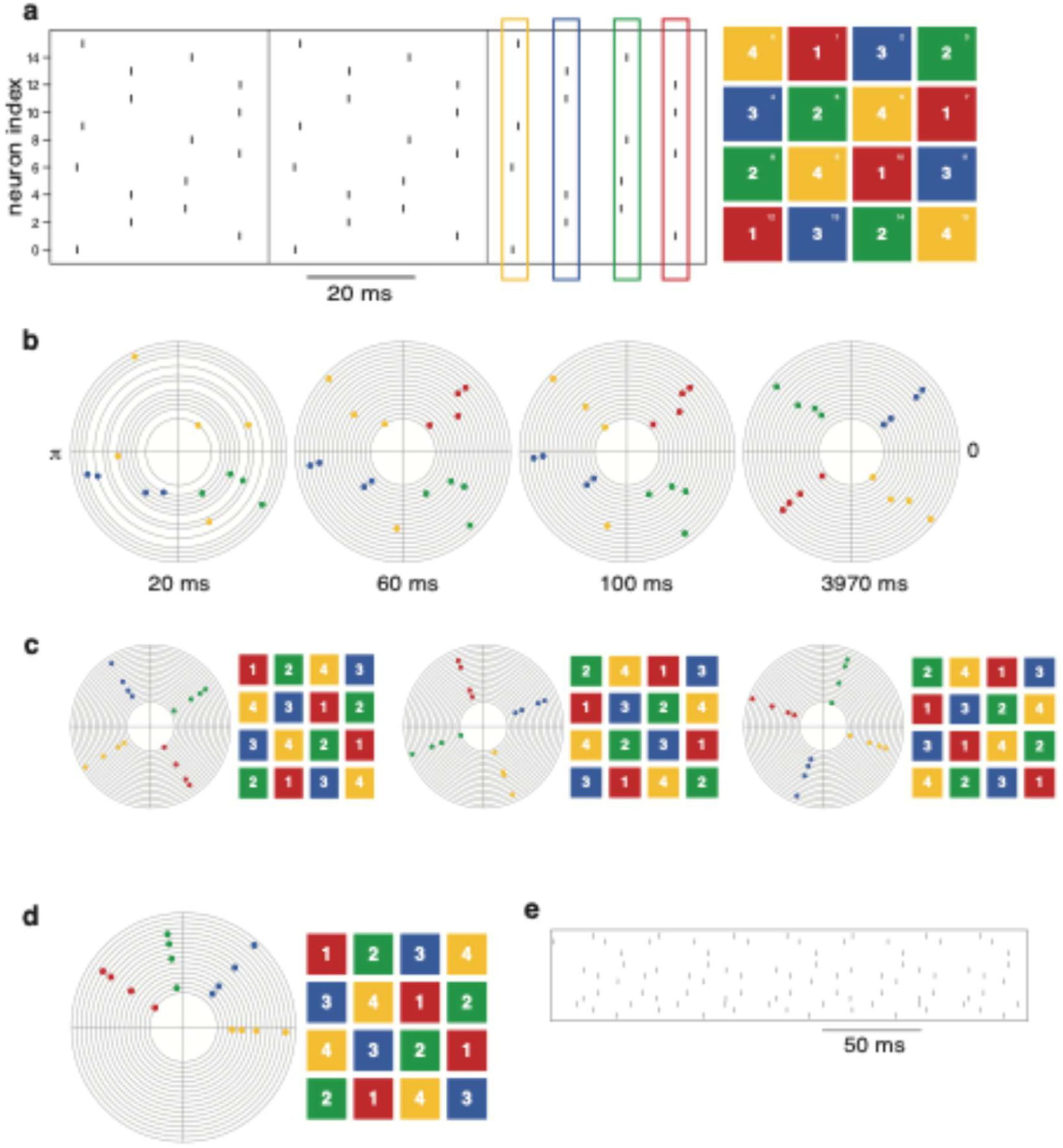
Oscillations plus depolarizing biases steer a 4×4 Sudoku network into specific solutions. (a) Raster (left) showing periodic spiking; corresponding Sudoku solution (right). (b) Network settling from random initial conditions without oscillation or biases; circular phase plots for cycles at labeled times. (c) Different random starts (no drive/bias) yield distinct steady states, each mapped to a Sudoku solution. (d) Global oscillation plus solution-specific depolarizing biases reliably produces the target Sudoku solution regardless of initial condition; the highest-bias group spikes first. (e) Biases alone (no oscillation) produce variable outcomes across initial conditions.

Each coloring of the Sudoku network partitions neurons into groups that are not inhibited by each other. Given one such coloring (a solution of a 4×4 Sudoku puzzle), we assigned a distinct depolarizing bias to each color (equivalently, digit from 1 to 4) group, so that all neurons of the same color spiked synchronously at a specific phase of the background oscillation. With four distinct biases applied to the four color groups, the network converged to a state in which each group spiked at a distinct phase of the oscillation, independent of the initial condition (**Fig. 3d**). In contrast, in the absence of the driving oscillation, the same four biases did not reliably drive the network to the same final state, in that the asymptotic activity depended on the initial condition (**Fig. 3e**). Thus, oscillations combined with appropriately chosen depolarizing biases can drive the network to one of many periodic steady states.

Next, we asked whether, in such a recurrent network, an oscillation together with a transient depolarizing bias could switch the system from one steady state to another. To this end, we first drove the network to a particular coloring (corresponding to a specific 4×4 Sudoku solution) using a set of depolarizing biases as described above, such that neurons of a given color had identical depolarizing currents. Once the network settled to its steady state spiking activity in the presence of solution-specific depolarizing biases, we removed the biases. For some colorings (each representing a 4×4 Sudoku solution), the network persisted in the same periodic state even after the biases were removed - we refer to these states as *stable periodic attractors* (**Fig. 4a**). For other colorings, though the network maintained its periodic spiking activity for the duration of the coloring-specific biases, once the biases were removed it transitioned into a new periodic steady state. In some instances, this new state corresponded to a different coloring (**Fig. 4b**). In other instances, the new state could be mapped to the same coloring, however, the order of synchronously spiking groups was permuted with respect to the original periodic state (**Fig. S2a**). For stable periodic attractors, once the dynamics had settled to its asymptotic state, we switched off the coloring-specific biases and used a transient perturbation consisting of depolarizing inputs designed to drive the system to a different stable attractor (**Fig. 4c**). The transient bias lasted for ∼10 cycles of the background oscillation. We found that during this transient window, the network’s spiking activity reliably switched from the first stable attractor state to the second and persisted in the second stable attractor state even after the input was removed (**Figs. 4c and S2b**). This behavior held true for all pairs of initial and target stable periodic attractor states that we simulated.

**Figure 4:**
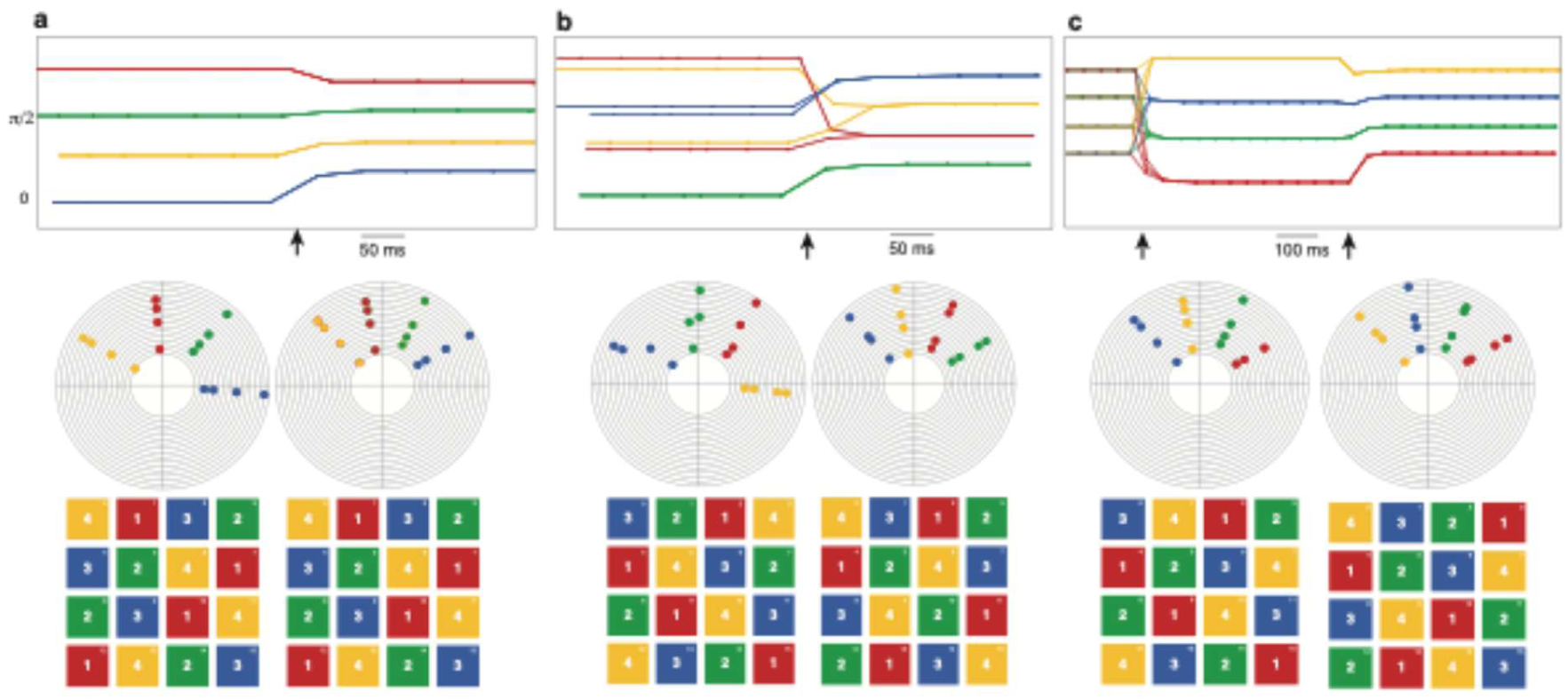
Transient perturbations switch periodic attractors. (a) Stable attractor: phase-time plot (top) and circular phase plots shown before and after turning off biases. The arrow indicates the time at which the bias was turned off. The phase ordering and the corresponding Sudoku solutions remain unchanged. (b) Unstable attractor: turning off biases causes synchronous groups to split and reassign, changing the coloring and Sudoku solution. (c) Switching attractors with a transient perturbation: applying a bias for 500 ms (between the two arrows) moves the network from coloring 1 to coloring 2, which persists after biases are removed; phase plots and Sudoku mappings show initial and final states.

In sum, a recurrent network can be reliably driven into specific periodic steady states by applying a coloring-specific set of depolarizing bias currents together with a common oscillatory background (**Fig. 3d**). A subset of these periodic attractors is stable, i.e., they persist even after the depolarizing bias currents are removed (**Fig. 4a**), and transient perturbations can switch the network between different stable attractors (**Figs. 4c and S2b**).

### Network asymmetries determine the stability of attractor states

A subset of colorings (or 4×4 Sudoku solutions) correspond to stable attractor states, in that they persist even when the depolarizing biases are turned off (**Fig. 4a**), while other colorings/Sudoku solutions do not (**Fig. 4b**). What gives rise to this differential stability? The collective dynamics of the neurons is characterized by two features, the *grouping*, viz., the identity of neurons that form a synchronized group, and the *order* in which these groups spike. The identity of neurons that spike together is determined by a coloring of the Sudoku network. But this does not impose any constraint on the ordering, which defines a permutation of the assigned colors. Yet, for a given coloring, some orderings are stable periodic attractors, but others are not (**Figs. 4a and S2a**).

To understand why some orderings are preferred, we considered a vastly simpler network, namely, a 9-partite network consisting of 9 groups of neurons (with nine neurons per group) with all-to-all inhibitory connections across groups and all-to-all excitation within each group (**Fig. 5a**). This network has a single 9-coloring, namely a unique color assigned to each of the 9 groups of neurons, analogous to the tripartite network in **Fig. 2a**. Neurons within a group spike together, while different groups spike at distinct times. Due to the network’s symmetry, no ordering is preferred, resulting in 9! potential orderings. The steady state ordering of the network depends on the initial condition (**Fig. S3a**). Analogous to the 4×4 Sudoku, when a background oscillation was broadcast to all neurons in combination with 9 distinct depolarizing biases to each of the 9 groups of neurons, the asymptotic ordering became independent of the initial condition (**Fig. S3b, c**). Specifically, the magnitude of the bias to each group determined the ordering. The group receiving maximum strength of depolarizing bias spiked first in every cycle, followed by the group receiving the second-highest depolarizing bias, and so on (**Fig. S3b**). Like the 4×4 Sudoku, the 9-partite network could also be switched between stable states, in this case different orderings, using transient perturbations (**Fig. S3d**). When we drove this symmetric 9-partite network to a given ordering using corresponding depolarizing biases and then switched off the biasing input once it reached steady state, we found that its ordering remained unchanged even after the bias was turned off (**Fig. 5b**). All orderings were stable periodic attractors.

**Figure 5:**
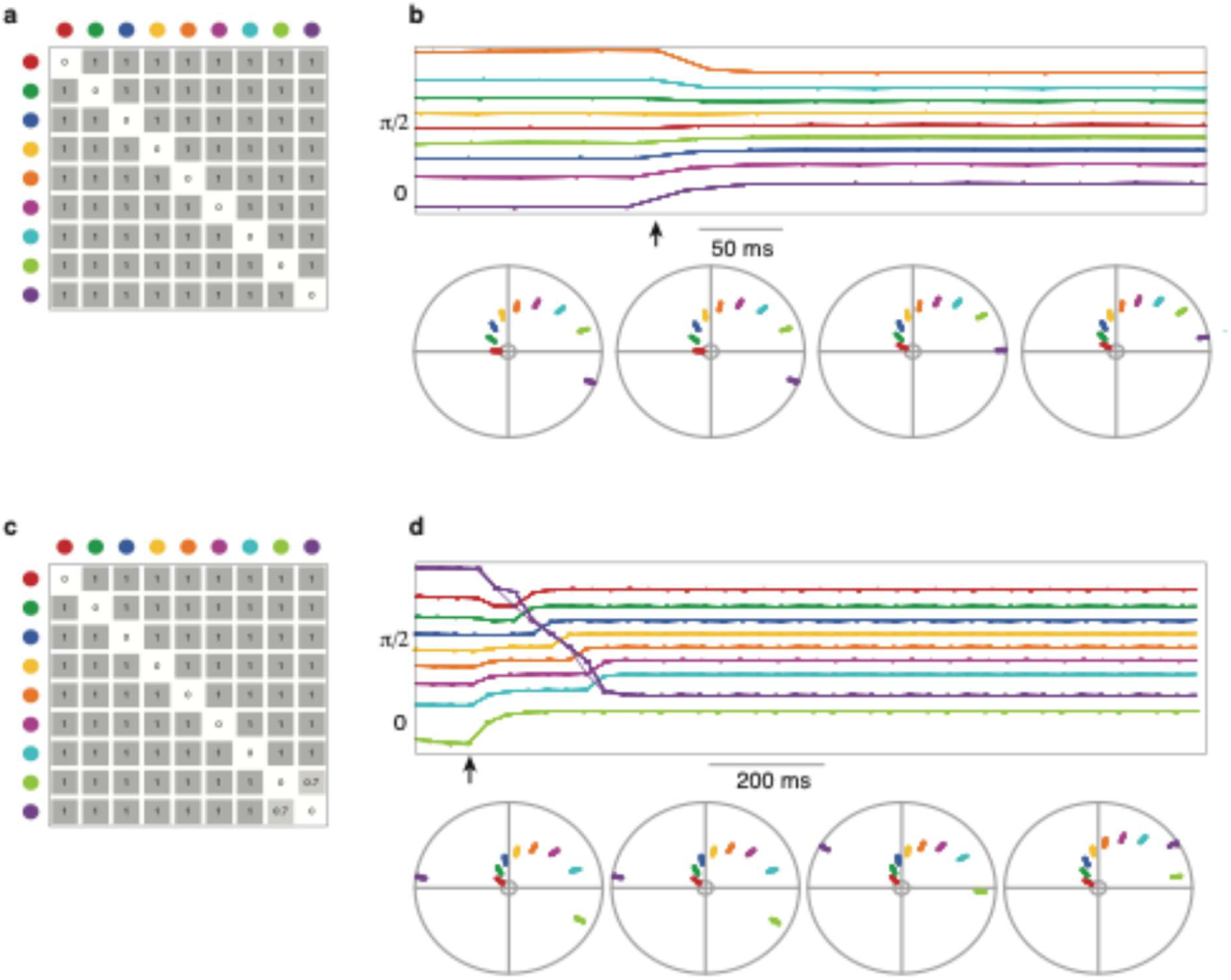
Inhibition strength sets ordering stability in a 9-partite network. (a) Symmetric connectivity matrix (groups of nine neurons; entry 1 = full inhibition). (b) With symmetry, all spike orderings remain stable before and after biases are removed (indicated by the arrow). The circular phase plots (bottom, left to right) show the phase of spiking changing progressively as the biases are removed (c) Connectivity with weakened inhibition between groups 8 and 9 (0.7). (d) Breaking symmetry makes orderings with groups 8 and 9 successive preferred; non-preferred orderings reconfigure to the favored ordering once biases are turned off.

Next, we systematically perturbed the network’s topology to break its symmetry, leading to the emergence of some preferred orderings. In particular, we reduced the strength of inhibition between the groups labeled 8 and 9 to 70% of the original strength while keeping all other coupling strengths unchanged (**Fig. 5c**). We expected that in this asymmetric network, orderings in which groups 8 and 9 spiked in succession would be favored due to reduced inhibition between them. We found that this was indeed the case - only states where groups 8 and 9 spiked in immediate succession were stable periodic attractors of the asymmetric network. Even though we were able to use a set of depolarizing biases to drive the network to a state where groups 8 and 9 did not spike in succession, once we removed the biasing input, the network quickly switched to a new ordering where groups 8 and 9 spiked in immediate succession (**Fig. 5d**). Thus, the topology of the asymmetric 9-partite network, particularly the strength of inhibition between pairs of groups, forced only a subset of states to be stable.

How can these observations be extended to the Sudoku network? After the depolarizing biases were removed, the 4×4 Sudoku network dynamics followed one of three possible outcomes: one, the network remained in the same periodic state (i.e., the state was a stable periodic attractor; **Fig. 4a**); two, it transitioned to a new periodic steady state corresponding to a different coloring (**Fig. 4b**); and three, it entered a new periodic state that mapped to the same coloring, but with a permuted order of synchronously spiking groups relative to the original state (**Fig. S2a**). From these observations, we identified general rules that determine whether a given set of depolarizing biases leads to a stable periodic attractor state, based on the symmetries of the state. Once a particular grouping and ordering of neurons is chosen and the corresponding depolarizing biases are used to drive the network, two conditions must be met for the resulting state to persist when the biases are turned off. First, for the grouping to be stable, every neuron within a group must receive the same number of inhibitory inputs from each of the other groups, so there are no within-group asymmetries in the network structure (**Fig. 6a** shows an example of within-group asymmetry). Second, for the ordering to be stable, groups that have fewer inhibitory connections between them must spike closer in time than groups that are more strongly connected by inhibition, since across-group asymmetries in inhibitory inputs render some temporal orderings stable and others unstable (**Fig. 6b** shows an example of across-group asymmetry).

**Figure 6:**
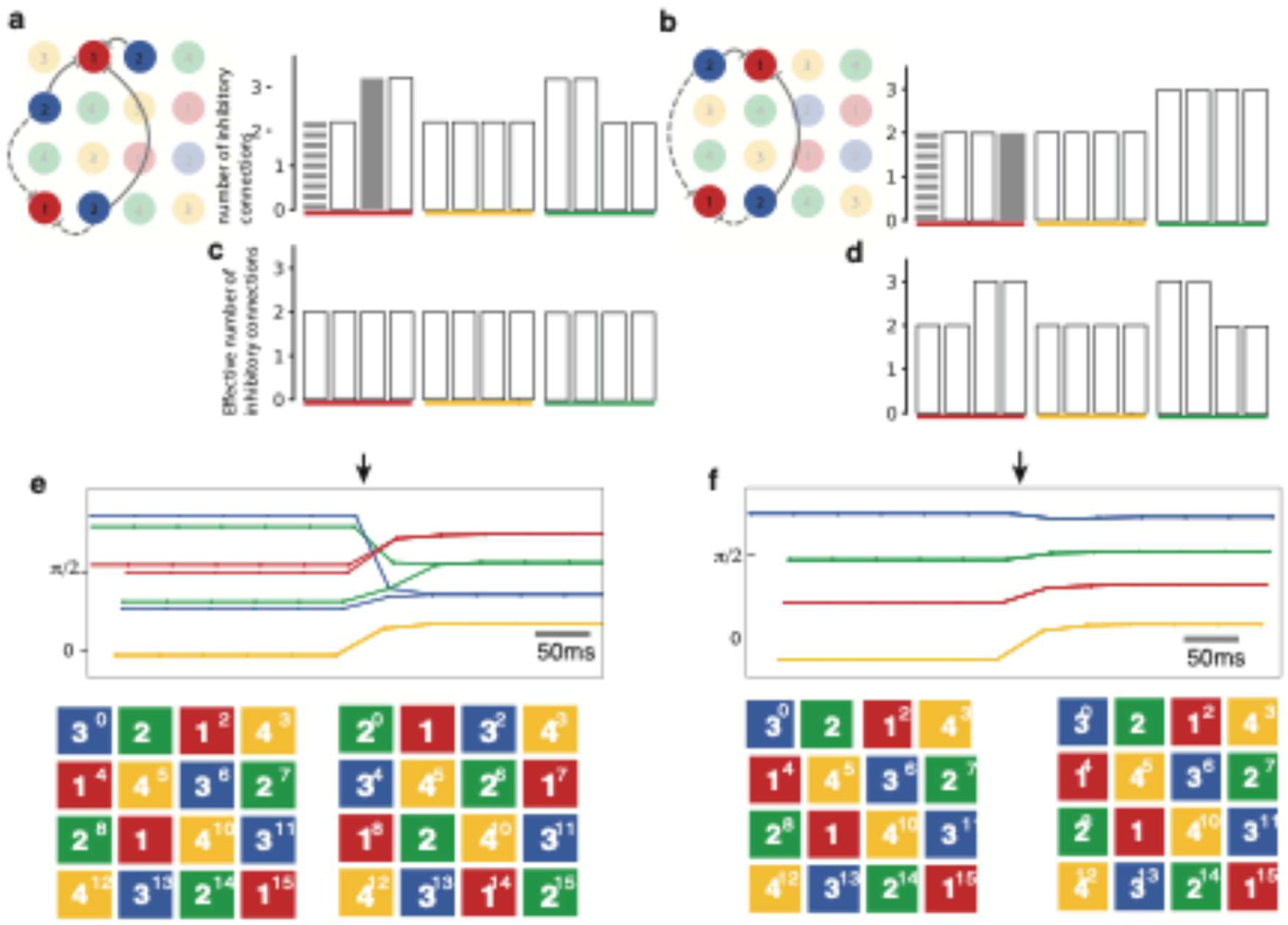
Stabilizing specific colorings by tuning the connectivity weights. (a) Sudoku solution with within-group inhibitory asymmetries, yielding an unstable coloring that changes when biases are removed; bar plot shows asymmetric inhibitory inputs from group 2 (blue) to groups 1, 3, and 4. The connections from group 2 to 1 are isolated in the network and highlighted in the corresponding bar plot. (b) Solution with only across-group asymmetries: coloring is stable but spike ordering is not; bar plot shows unequal inhibitory inputs from group 2 (blue) to groups 3 and 4. (c) Coloring-specific synaptic weight normalization equalizes effective inhibitory inputs (2 per neuron) across and within groups for that coloring, removing asymmetries; excitatory weights are similarly normalized (not shown). (d) The same normalization does not, in general, remove asymmetries for a different coloring, leaving within- and across-group heterogeneity in inhibitory inputs. (e) Before normalization, colorings with within-group asymmetries are unstable once biases are turned off. (f) After normalization, both the coloring and ordering become stable and persist without biases.

Thus, we show that the symmetries of a given state, defined with respect to the underlying network topology, govern the stability of that state. Furthermore, we establish well-defined rules for predicting the stability of any given state using its symmetries.

### Individual colorings can be stabilized by tuning the weights of network connectivity

In the 9-partite network, all orderings were stable when the network connectivity was symmetric (**Fig. 5a, b**), but when we broke the symmetry by changing the weights of network connectivity, some orderings became unstable (**Fig. 5c, d**). That is, the stability of a particular state could be changed by altering the weights of connections between specific neurons in the network. Applying the same idea to the 4×4 Sudoku network, we asked if unstable states of the network could be converted into stable periodic attractor states by tuning the excitatory and inhibitory connection weights.

As we saw previously, one of the conditions for a state to be stable is that each neuron within a group must receive identical total inhibition from every other group. When this condition does not hold (**Fig. 6a**), the corresponding grouping (coloring) is unstable, i.e., it persists while the depolarizing biases are on, but switches to a new grouping (coloring) once the biases are turned off (**Figs. 4b and 6e**). To make a given coloring (corresponding to a Sudoku solution) satisfy this condition, we employed a normalization strategy to tune the weights of the network. Specifically, for each neuron, we counted the number of inhibitory and excitatory connections that it received from a particular group and normalized the weights of all connections from that group to that neuron, such that it now received the equivalent of two inhibitory inputs and one excitatory input from that group. We repeated this process for all 16 neurons in the 4×4 Sudoku network. The modified weight matrix ensured that for the given coloring (Sudoku solution), each neuron within a group now received identical total inhibition from every other group (**Fig. 6c**). It is important to note that the normalization process was specific to a given coloring, and therefore normalization did not ensure that the condition was satisfied for other colorings (**Fig. 6d**). Once the network weights were normalized in this manner for a given coloring, the coloring became fully stable, i.e., the grouping persisted even after removal of the depolarizing biases (**Fig. 6f**). Furthermore, the normalization ensured that the total inhibition strength between all pairs of groups became equal, and therefore all orderings became stable, i.e., there was no preferred ordering (**Fig. S4**). Thus, the weights of the network could be tuned in a coloring-specific manner to ensure that the coloring, as well as all orderings corresponding to that coloring became stable.

### Asymmetries along with crowding affect network dynamics at higher group number

A 2^2^×2^2^ (4×4) Sudoku has 288 possible solutions. Increasing the size of the Sudoku to the more commonly encountered 3^2^×3^2^ (9×9) grid (e.g., **Fig. 7a, b**) leads to a combinatorial explosion in the number of possible solutions (∼10^9^)^11^. We now have 9 clusters of neurons spiking at distinct phases of the oscillation. As the number of groups increases, the phase difference between groups decreases. Further, within-group asymmetries can result in neurons within a group spiking at slightly different phases instead of being perfectly synchronous. We asked if these competing influences, i.e., phase variability due to asymmetries and crowding due to an increased number of synchronized groups, destabilizes some solutions of the 9×9 Sudoku.

**Figure 7:**
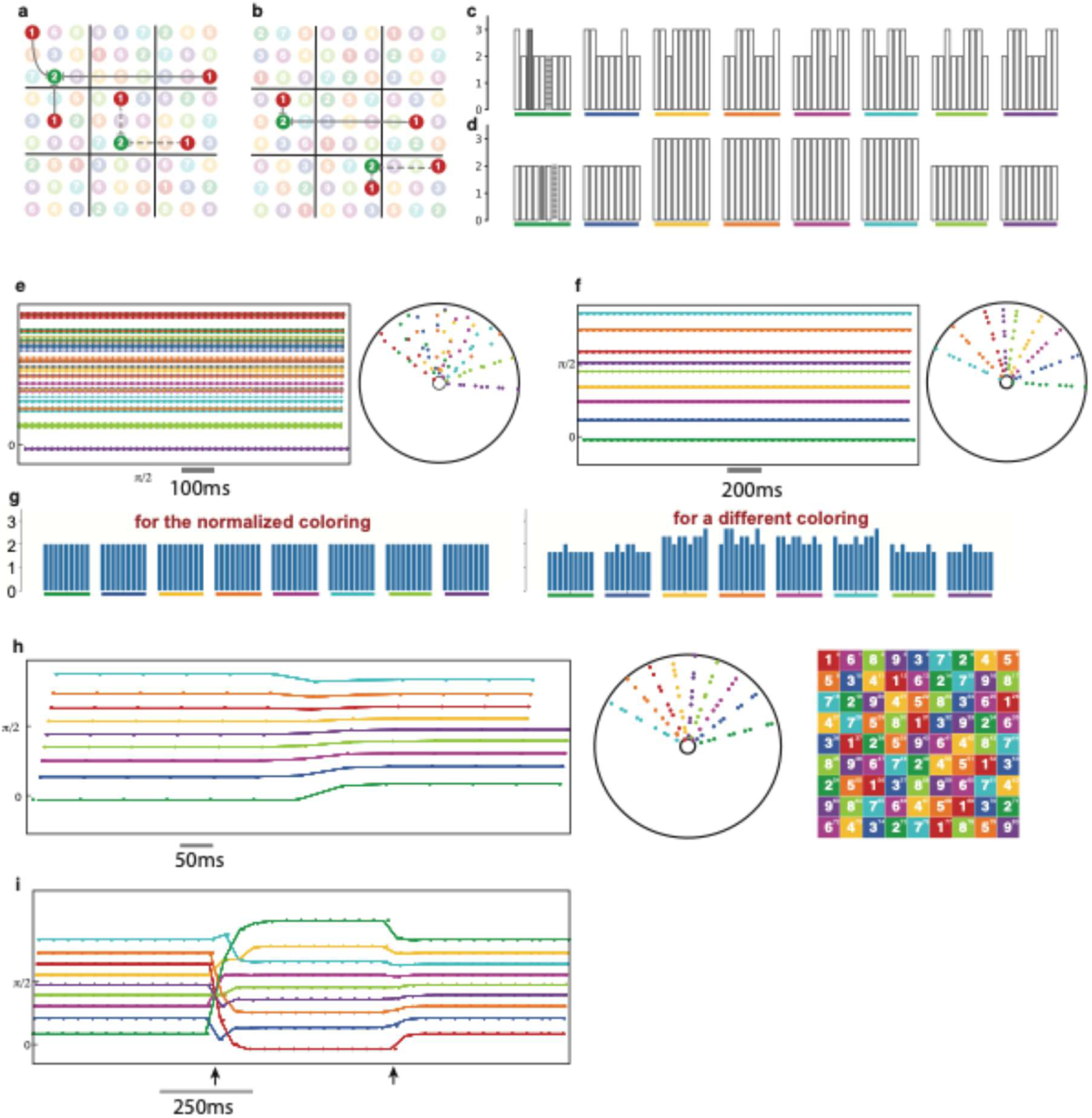
Asymmetries and crowding shape dynamics in a 9×9 Sudoku network. (a) Sudoku solution with within-group inhibitory asymmetries; bar plot (c) shows unequal inputs from group 1 (red) to neurons in groups 2–9, marked by filled/dashed bars corresponding to example connections. (b) Solution with only across-group asymmetries; bar plot (d) shows differences in inputs across groups, e.g., 3 (blue) vs. 4 (yellow), 7 (cyan) vs. 8 (green). (e) With a common oscillation and biases for a coloring with within-group asymmetries, the steady state lacks 9 distinct synchronous groups and cannot be mapped to a valid Sudoku solution. (f) With oscillation and biases for a coloring with only across-group asymmetries, some initial conditions yield valid Sudoku solutions, but the spiking order need not align with bias order. (g) Coloring-specific weight normalization equalizes effective inhibitory inputs (2 per neuron) and excitatory inputs (1 per neuron, not shown), removing both within- and across-group asymmetries for that coloring; for a different coloring, asymmetries persist. (h) After normalization, all orderings of that coloring become stable periodic attractors: spiking order matches bias order and persists after biases are removed, as shown by the circular phase plot and Sudoku mapping. (i) A transient perturbation (brief application of biases for a new ordering) can switch the network from one stable ordering to another, which then remains stable after biases are turned off.

To answer this question, we simulated a 9×9 Sudoku network analogous to the 4×4 Sudoku network. Each neuron received a background oscillatory drive, and a depolarizing bias. Like in the 4×4 Sudoku, for a given coloring (Sudoku solution) and ordering of neurons, we assigned a distinct depolarizing bias to each color (or equivalently, digit from 1 to 9) to drive the network to that state. We compared the network dynamics for colorings that showed within-group asymmetries (**Fig. 7a, c**) to those that did not (**Fig. 7b, d**). When we drove the network to a coloring (representing a 9×9 Sudoku solution) having within-group asymmetries, we found that the neurons within groups were not perfectly synchronous and instead spiked at slightly different phases. Because of this, and due to increased crowding in the 9×9 Sudoku network, the different groups merged into each other, and the periodic spiking activity of the network could not be mapped to a Sudoku solution (**Fig. 7e**). On the other hand, when we drove the network to a coloring that did not have within-group asymmetries but had across-group asymmetries, the network’s steady state spiking activity matched the coloring, i.e., it mapped to the given Sudoku solution for some initial conditions but not for others. However, unlike in the case of the 4×4 Sudoku network, even when the coloring mapped to the solution of the 9×9 Sudoku the ordering did not always match the relative magnitudes of depolarizing biases, e.g., the group with the 4^th^-largest magnitude of depolarizing bias was not always the 4^th^ group to spike at steady state (**Fig. 7f**).

Next, we asked if any given coloring (9×9 Sudoku solution) and ordering could be made reliably accessible using a corresponding set of depolarizing biases, by modifying the weights of network connectivity. To test this, we normalized the weights of inhibitory and excitatory inputs to each neuron following the same approach used for the 4×4 Sudoku network, i.e., for the given coloring (solution), the weights were tuned such that each neuron of a group received the equivalent of two inhibitory inputs (**Fig. 7g, left**) and one excitatory input from every other group. The normalization was coloring-specific, therefore in general it did not eliminate asymmetries for other colorings/solutions of the network (**Fig. 7g, right**). When within and across-group asymmetries were eliminated by this method for a given solution, that solution and all its orderings became stable periodic attractors, i.e., they could be accessed by the corresponding depolarizing biases independent of the initial condition and persisted even after the biases were removed (**Fig. 7h**). Further, a transient perturbation could be used to switch between different orderings of the given coloring/solution (**Fig. 7i**).

In sum, we found that asymmetries and crowding effects influence the stability of solutions in larger Sudoku networks. By normalizing input weights to eliminate these asymmetries, we achieved stable solutions in the 9×9 Sudoku network, enabling reliable transitions between different orderings through transient perturbations.

## Discussion

Hopfield and colleagues^8,13^ showed that phase-of-firing codes can generate concentration-invariant representations of odor stimuli in models of the olfactory system where neurons receive rhythmic drive from respiration. Notably, these models lack lateral or recurrent connectivity. In such feedforward architectures, temporal information is referenced to a common oscillation without being strongly shaped by network dynamics. However, many brain regions in which temporal coding is believed to be critical, including hippocampus^4,14^, movement circuits^15^, and olfactory networks^16–19^, exhibit dense recurrent connectivity^20,21^, and in these systems, temporally ordered activity patterns are thought to play a central role in functions such as memory retrieval, motor planning^22^, and sensory discrimination^23^.

In this context, our results extend the phase-of-firing code beyond purely feedforward layers to recurrently connected networks. By introducing structured recurrent connectivity, we demonstrate how phase-of-firing codes can be both implemented and stabilized in recurrent circuits, thereby broadening their relevance from the sensory periphery to cortical and subcortical networks. Moreover, the networks we study express intrinsic oscillatory activity, akin to theta or gamma rhythms^24^, which shape network dynamics. Our model shows that such oscillations can be exploited to selectively stabilize and control temporal sequences, providing a mechanistic account of how phase-of-firing codes might operate within biologically realistic recurrent network architectures.

From the perspective of associative memory, our framework differs in important ways from classical Hopfield networks. Traditional Hopfield models, built on Hebbian learning, are limited to storing on the order of 0.13-0.14 patterns per neuron, and their attractors are fixed points corresponding to static patterns^25^. In contrast, the oscillator network examined here supports a rich repertoire of patterns, with capacity modulated by the balance between excitation and inhibition^12^. The stable states in our model take the form of periodic sequences rather than static configurations, so information is encoded both in the identity of active neurons and in their temporal ordering across cycles of the background oscillation.

This additional temporal dimension permits a substantially larger number of distinguishable patterns than in standard Hopfield networks. At the same time, the combinatorial explosion of possible periodic states renders an exhaustive characterization of basins of attraction infeasible. We address this by showing that a common oscillatory drive along with specific depolarizing biases can reliably select particular network states, and that brief, pattern-specific inputs can flexibly transition the system between distinct periodic attractors. Importantly, the persistence of a given state after inputs are removed depends on its symmetry properties: network states whose groupings and orderings are aligned with structural symmetries are preferentially stabilized, whereas asymmetric states tend either to reorganize or to collapse into more symmetric configurations. Thus, the symmetry structure of the underlying connectivity provides predictive constraints on which dynamic memories can be stably expressed.

Our findings also speak to the role of active sensing in shaping temporal codes for odor. Animals do not sample odors passively; instead, they adjust sniffing rate as a function of odor concentration, novelty, and decision demands, so that the spatiotemporal representation of an odor unfolds across multiple sniff cycles^26^. In insects, analogous active sampling arises as rhythmic wing beats drive periodic airflow over the antennae, structuring when odor molecules reach receptor neurons^27^. Within this actively controlled sampling regime, a phase-of-firing code offers a natural mechanism for representing odor identity in a way that is relatively invariant to changes in sniffing or wing-beat frequency.

Because spikes are referenced to the phase of an ongoing oscillation, modulations in sampling frequency primarily reshape the temporal scaffold rather than the relative phase relationships that carry stimulus identity. Additional behaviors that intermittently perturb the sensory stream, such as antennal flicks, may further enrich the code by rapidly exposing features of the odorant that would otherwise emerge only slowly over many regular cycles. Our model provides a dynamical systems perspective on these processes: transient perturbations to the input switch the network between distinct periodic attractors, effectively routing activity through different regions of phase space. Viewed in this way, active sensing behaviors can be interpreted as controlled perturbations that enable the system to explore its repertoire of odor-evoked trajectories more thoroughly, enhancing both the richness and flexibility of the temporal code implemented by recurrent neural circuits.

## Methods

### Neuronal network simulations using Brian2

Conventional integrate-and-fire neurons were used for all simulations, using the Brian2 module in Python^28^. The following set of equations were specified for each neuron.

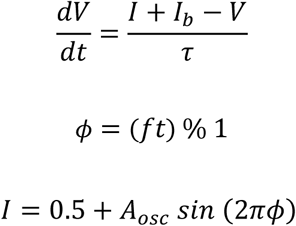

where *V* is the neuron’s potential, *τ* is a time constant, *I* is an oscillatory input current, *f* is the frequency of the oscillatory input, *ɸ* is the phase of the oscillatory input (between 0 and 1), *A_osc_* is its amplitude, and *I_b_* is an input bias current (for coloring-specific depolarizing biases), which was specified separately for each neuron.

For all simulations, the value of time constant used was *τ* = 10 ms. The value of frequency was *f* = 30 Hz for the 9-partite networks, and *f* = 25 Hz for the Sudoku networks. *A_osc_* was 0.2 for all simulations, and the range of values for *I_b_* was optimised by the method described in a later section. The default timestep of 0.1 ms was used for all simulations.

In addition to these equations, synaptic connections were defined according to an input connectivity matrix corresponding to the network being simulated, namely 4×4 Sudoku, 9×9 Sudoku or 9-partite networks (see next section). There was no time delay of transmission. Constant strengths of inhibition (*C_inh_*) and excitation (*C_exc_*) were defined for all connections. In some simulations (where a tuning of synaptic weights was performed), a relative weight of synaptic connections was defined to modulate the strength of individual synapses.

The Synapses object of the Brian2 module^28^ was used to define the synaptic connections and strength. Spike timings of each neuron were recorded using the SpikeMonitor object, and when an oscillatory input current was present, the value of *ɸ* at every firing was also recorded for each neuron. The StateMonitor object was used to record time series of the potential *V* for all neurons, and to record the oscillatory input. Random initial conditions were chosen at every run, i.e., each neuron got a random initial value between 0 and 1 of its potential *V*.

### Network connectivity matrices

The binary connectivity matrix for a 9-partite network was constructed as an 81×81 matrix, with each group having 9 neurons. Each neuron in a particular group had an inhibitory connection to every other neuron that did not belong to its group (see **Figs. 2b and 5a**). Excitation was complementary to inhibition, therefore all neurons within a group had excitatory connections between them.

In the Sudoku networks (both 4×4 and 9×9), the inhibitory connections were defined by the rules of Sudoku. Each “cell” of the Sudoku was represented by a single neuron. Thus the 4×4 and 9×9 Sudokus had connectivity matrices of dimension 16×16 (see **Fig. 2d**) and 81×81 respectively. Each neuron had an inhibitory connection to all the other neurons in the same row, same column and same smaller square (2×2 in case of the 4×4 Sudoku and 3×3 in case of the 9×9 Sudoku) of the Sudoku grid. Again, excitation was complementary to inhibition i.e. any pair of neurons which did not have an inhibitory connection were connected by an excitatory connection.

### Tuning of synaptic parameters

The synaptic strengths of excitation (*C_exc_*) and inhibition (*C_inh_*) were tuned by trial and error, for all the networks used (9-partite, 4×4 Sudoku and 9×9 Sudoku). To do this, *I_b_* was first fixed as a constant value for all neurons, and the strengths *C_exc_* and *C_inh_* were tuned such that each neuron fired exactly once in a cycle of the driving oscillation. This process was repeated for various fixed values of *I_b_*, and the parameters (a combination of *I_b_*, *C_exc_* and *C_inh_*) which produced the most repeatable periodicity across trials with random initial conditions were chosen for further simulations of that network. For the 9-partite network, *C_exc_* = 0.003 and *C_inh_* = −0.005. For the 4×4 Sudoku, *C_exc_* = 0.0028 and *C_inh_* = −0.05. For the 9×9 Sudoku, *C_exc_* = 0.0028 and *C_inh_* = −0.027. When coloring- or ordering-specific depolarizing biases had to be provided, *I_b_* was adjusted for each neuron individually as described in the next section.

### Finding optimal range of input bias currents

To find the optimal range of input bias currents *I_b_* for which the firing of neurons would be periodic with the driving oscillation, a raster plot was made in which bias current increased with increasing neuron index along the y-axis (**Fig. S1**). This was similar to Fig. 1 of Brody and Hopfield, 2003^8^. There was a clear range of values of the bias current for which the firing was periodic with the driving oscillation, and each neuron fired exactly once per cycle. In this range, each neuron fired at exactly the same phase (*ɸ* value) of the driving oscillation in every cycle, i.e., it was phase-locked. The phase at which a particular neuron fired increased linearly with the bias current over this range: the higher the bias current, the earlier it fired in the cycle of the oscillation. This range of *I_b_* provided a first estimate for the value to be used in the simulations. However, since this range was determined using a set of neurons with no network connections (excitatory or inhibitory), it had to be fine tuned (typically shifted to a slightly higher range) to get periodic phase-locked firing in the simulations of Sudoku and 9-partite networks (which have recurrent connectivity between neurons).

## Acknowledgements

We thank members of Assisi and Nadkarni labs at IISER Pune for useful discussions. We also acknowledge the National Supercomputing Mission (NSM) for providing the computing resources of ‘PARAM Brahma’ at IISER Pune, which is implemented by C-DAC and supported by the Ministry of Electronics and Information Technology (MeitY) and Department of Science and Technology (DST), Government of India.

## Data and Code Availability

All code, and raw data used to generate the figures is available upon request.

## Supplementary figures

**Figure S1:**
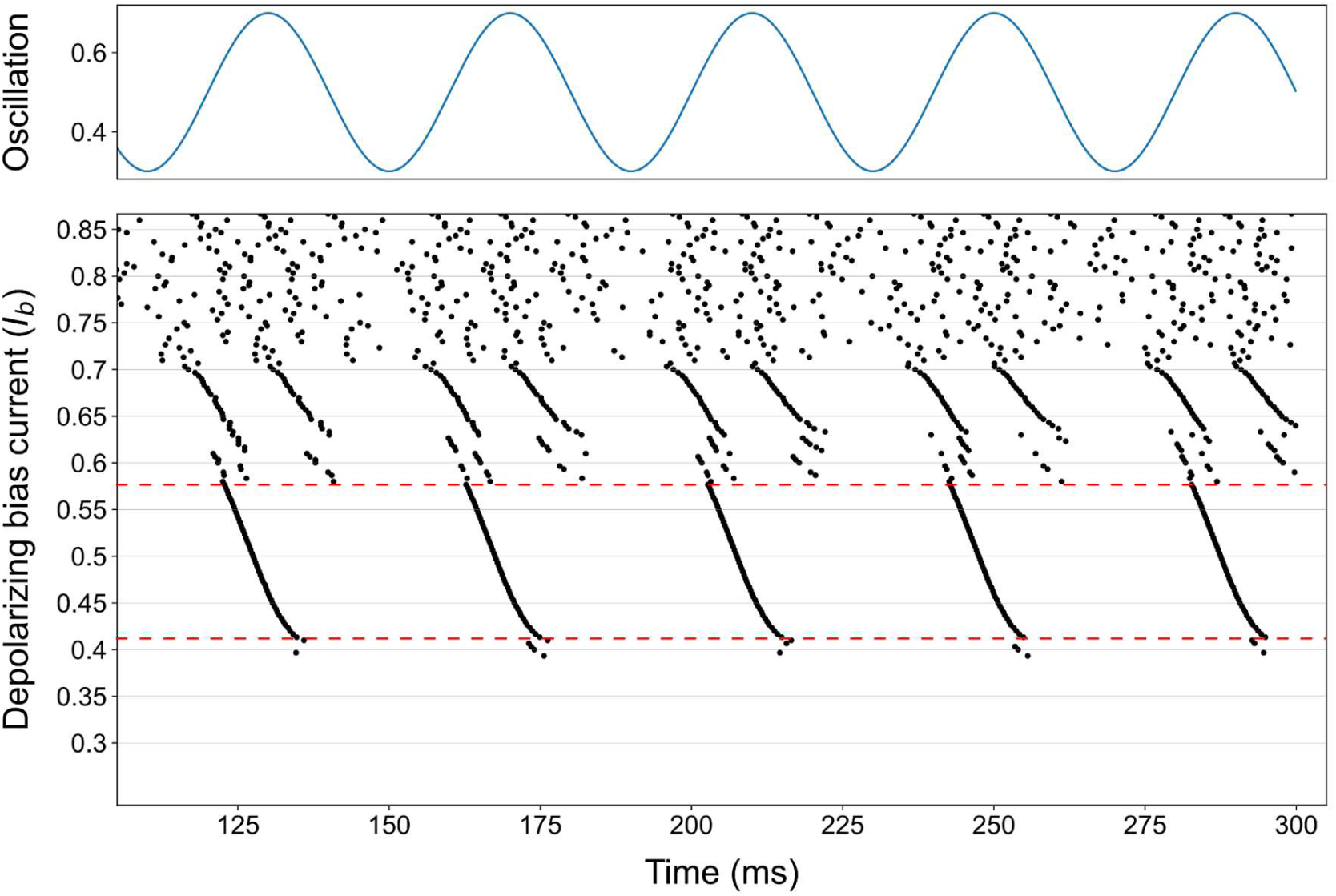
Determining range of depolarizing biases for which phase of firing (spiking) varies monotonically with magnitude of bias. Raster plot of uncoupled neurons driven by a common oscillation, each with a distinct depolarizing bias (y-axis increasing bottom to top). Red dotted lines mark the bias range in which each neuron fires exactly once per cycle and its spike phase varies monotonically with bias magnitude.

**Figure S2:**
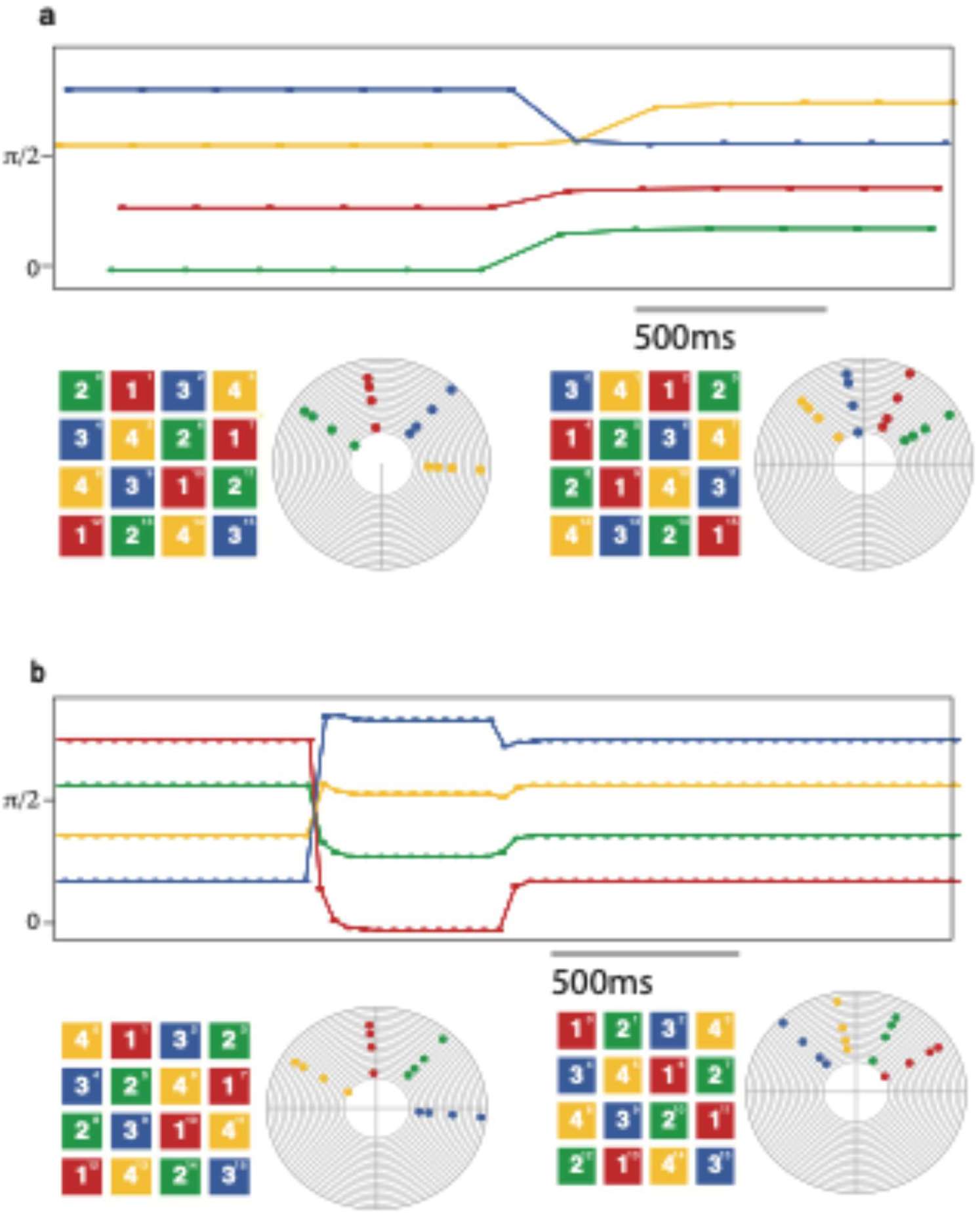
Transient perturbations switch spike order within a fixed coloring. (a) Unstable ordering: when depolarizing biases are turned off, synchronous groups keep the same coloring but change their spiking order (e.g. blue and yellow swap), yielding a permuted color assignment on the Sudoku grids. (b) Stable-to-stable switch: a brief application of biases for ordering 2 transforms the phase-lock pattern from ordering 1 to ordering 2, which then persists after biases are removed; circular phase plots and Sudoku grids show the initial and final orderings.

**Figure S3:**
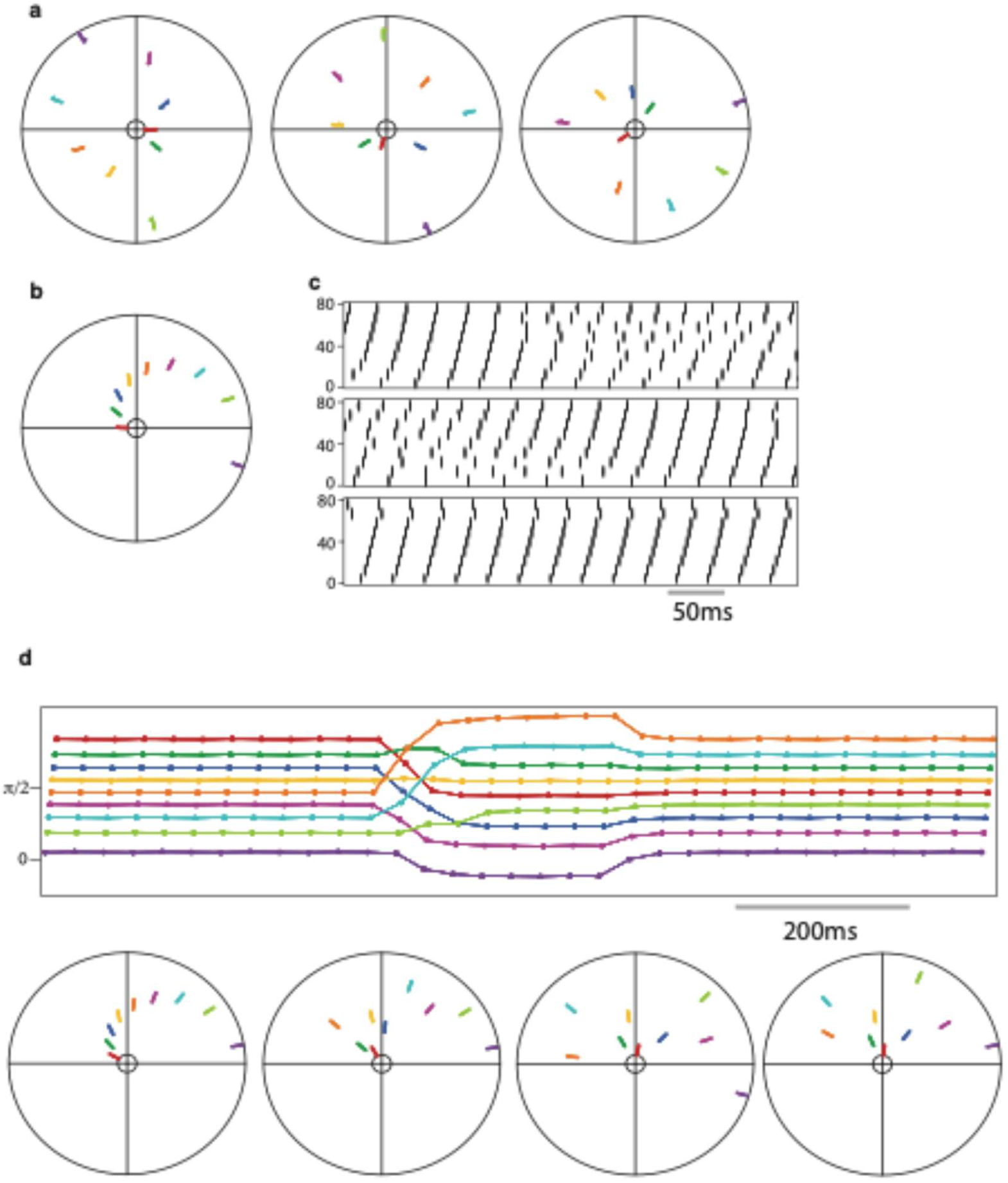
Oscillations and depolarizing biases control spike ordering in a 9-partite network. (a) Without oscillation or biases, random initial conditions lead to different steady-state spike orderings across trials. (b) Adding a common oscillation plus ordering-specific depolarizing biases drives the network reliably into the corresponding ordering, with the highest-biased group spiking first. (c) Biases alone, without oscillation, yield variable dynamics: steady states are not always periodic and their spike order may not match the bias order. (d) A brief application of biases for ordering 2 switches the network from a stable ordering 1 to stable ordering 2, which then persists after biases are removed.

**Figure S4:**
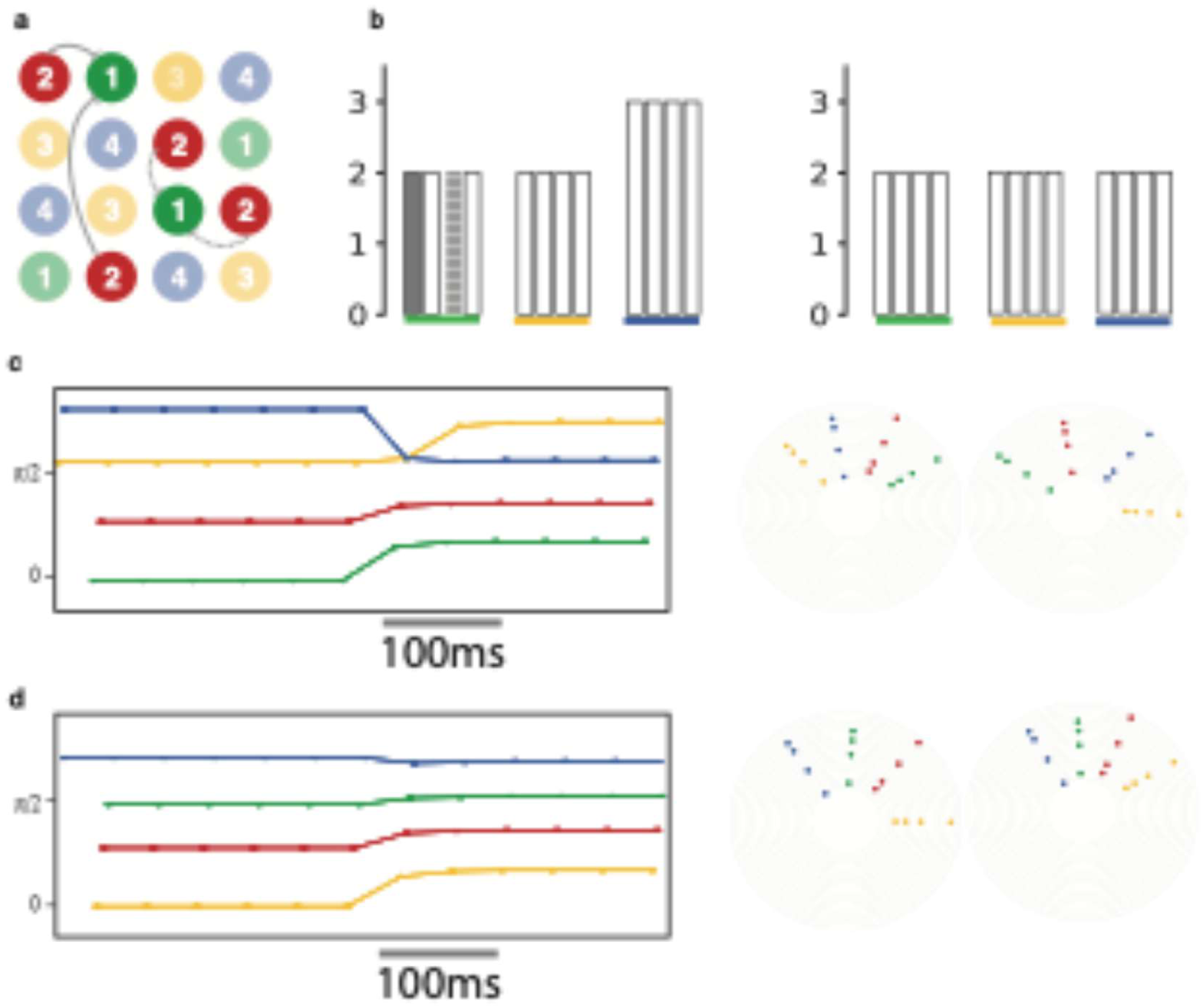
Weight normalization stabilizes all orderings of a given coloring. (a) Sudoku solution with across-group inhibitory asymmetries; bar plot shows unequal inputs from group 2 (red) to neurons in groups 1, 3, and 4. (b) Coloring-specific synaptic weight normalization equalizes effective inhibitory inputs (2 per neuron) for that coloring, removing across-group asymmetries; excitatory inputs are similarly normalized (not shown). (c) Before normalization, some spike orderings for this coloring are unstable and change once depolarizing biases are removed. (d) After normalization, all orderings of the coloring become stable and persist when biases are turned off.

